# Inhibition of SARS-CoV-2 infection by *Porphyromonas gingivalis* and the oral microbiome

**DOI:** 10.1101/2024.02.27.582258

**Authors:** Alexander Bontempo, Alexandra Chirino, Alireza Heidari, Alexandra Lugo, Satoru Shindo, Maria R. Pastore, Sibel A. Antonson, Cristina Godoy, Frank C. Nichols, Jan Potempa, Mary Ellen Davey, Toshihisa Kawai, Mark J. Cayabyab

## Abstract

The COVID-19 pandemic persists despite the availability of vaccines, and it is therefore crucial to develop new therapeutic and preventive approaches. In this study, we investigated the potential role of the oral microbiome in SARS-CoV-2 infection. Using an *in vitro* SARS-CoV-2 pseudovirus infection assay, we found a potent inhibitory effect exerted by *Porphyromonas gingivalis* on SARS-CoV-2 infection mediated by known *P. gingivalis* compounds such as phosphoglycerol dihydroceramide (PGDHC) and gingipains as well as by unknown bacterial factors. We found that the gingipain-mediated inhibition of infection is likely due to cytotoxicity, while PGDHC inhibited virus infection by an unknown mechanism. Unidentified factors present in *P. gingivalis* supernatant inhibited SARS-CoV-2 likely via the fusion step of the virus life cycle. We addressed the role of other oral bacteria and found certain periodontal pathogens capable of inhibiting SARS-CoV-2 pseudovirus infection by inducing cytotoxicity on target cells. In the human oral cavity, we observed the modulatory activity of oral microbial communities varied among individuals in that some saliva-based cultures were capable of inhibiting while others were enhancing infection. These findings contribute to our understanding of the complex relationship between the oral microbiome and viral infections, offering potential avenues for innovative therapeutic strategies in combating COVID-19.

## Introduction

As of August 2023, WHO estimated 770 million cases and 7 million deaths due to COVID-19 around the world. COVID-19 remains a major global public health problem despite the availability of effective vaccines. This can be attributed largely to the continuous emergence of SARS-CoV-2 variants and socioeconomic factors including vaccine hesitancy and lack of access to healthcare.^1,2^ It is therefore critically necessary to constantly find new avenues to treat and prevent infection.

The human mouth contains a tremendously large and diverse microbiota that is second only to the gut, with over 700 species of resident bacteria. Evidence continues to accrue supporting the essential role that the oral and human microbiome plays in health and disease,^3,4^ including COVID-19.^5-8^ A number of studies have attempted to establish an association between the oral and gastrointestinal microbiome with COVID-19 as well as other viral infection.^5,8-10^ For example, microbiome dysbiosis in both gut and mouth as result of an altered microbial composition was associated with a more severe COVID-19 infection.^8,10^ In addition, people with periodontitis were found to be more likely to have severe disease or die from COVID-19.^11-13^ Collectively, these studies strongly suggests that oral microbiome dysbiosis or potentially specific oral pathogens that are associated with periodontitis could play a role in COVID-19 infection and disease severity.

It is speculated that the oral microbiome could impact SARS CoV-2 infection through its interaction with the host. The oral microbiome has been shown previously to modulate innate and adaptive immunity, which could exacerbate SARS CoV-2 infection and disease.^14,15^ Lipopolysaccharides (LPS)-producing bacteria found in the oral cavity of COVID-19 patients exert an inflammatory effect.^16^ Oral bacteria species such as *Prevotella* and *Veillonella*, which are also commonly found in the oral cavity of COVID-19 patients may stimulate the production of IL-6, IL-23 and IL-1 inducing unwanted inflammation.^17^ There is evidence to suggest that in people with respiratory conditions such as pneumonia and chronic obstructive pulmonary disease, translocation of periodontal pathogens to the lungs may facilitate SARS CoV-2 replication in lung cells.^18,19^

Alternatively, oral microbes could directly interact with SARS CoV-2 and modulate infection. The potential for oral bacteria to contain anti-COVID-19 properties is possible since bacteria are known to possess antiviral compounds including LPS, polyketones, alkaloids, peptides, polyphenols, pyrones, quinones, sterols and terpenoids.^20,21^ Depending on the chemical properties of the antiviral compound and the specific virus, the mechanisms of viral inhibition have been shown to target virtually all the stages of the viral life cycle such as viral attachment and penetration, genome integration, replication and transcription.^20^

In the present study, we began to address the role of the oral microbiome in COVID-19 infection. We assessed the role of *Porphyromonas gingivalis* based on the observation that patients with periodontitis, which is an oral disease associated with *P. gingivalis* infection, had a higher risk of complications from SARS CoV-2 infection.^3,22,23,24^ A previous study showed *P. gingivalis* LPS was able to induce the expression the SARS-CoV-2 receptor ACE2 and the accessory protease TMPRSS2 in human gingival fibroblasts,^23^ suggesting a possible mechanism that could explain the increased severity of CoV-2 infection among periodontitis patients.

We also investigated the possible impact of the oral salivary microbiome from healthy human volunteers as well as laboratory strains of various commensal and pathogenic oral bacteria on SARS-CoV-2 infection. Important results of these studies are described herein showing that certain known compounds as well as unidentified factors are capable of markedly inhibiting SARS-CoV-2 infection in vitro. Interestingly, oral bacterial communities cultured from healthy human volunteers exhibited variable modulatory activities towards SARS-CoV-2. Altogether, these findings suggest a possible role for the oral microbiome in COVID-19 infection and further studies are needed to validate these observations *in vivo* or in the clinical setting.

## Results

### *P. gingivalis* factors inhibited SARS-CoV-2 pseudoviral infection

Certain factors produced by *P. gingivalis* have previously been shown to have antiviral properties.^25,26^ We tested whether *P. gingivalis* contained anti-SARS-CoV-2 properties and found that bacterial factors present in *P. gingivalis* supernatants markedly inhibited lentiviral vectors pseudotyped with SARS-CoV-2 spike protein (Fig. 1a). Infection of ACE2+ 293T cells with pseudoviruses containing either spike (S) protein cloned from either the original Wuhan strain or the Omicron BA.4/5 variant was significantly inhibited (Fig. 1a and b). The antiviral effect observed was specifically mediated by SARS-CoV-2 spike proteins in that VSV-G pseudotyped viruses were unaffected (Fig. 1c). The anti-SARS-CoV-2 effect exerted by *P. gingivalis* supernatant was substantial with an IC50 of 0.3% v/v. Using the WST-1 assay, we found that higher concentrations (≥7.5% v/v) were cytotoxic to the target cells (Fig. 4c). *P. gingivalis* supernatants produced with 10% FBS DMEM and BHI cultures were significantly more inhibitory than those produced with TSB. In addition to *P. gingivalis* supernatants, we also tested *P. gingivalis* lysates for inhibition and found that *P. gingivalis* lysates were highly cytotoxic even at low concentrations (less than 5%) (data not shown).

**Fig. 1.**
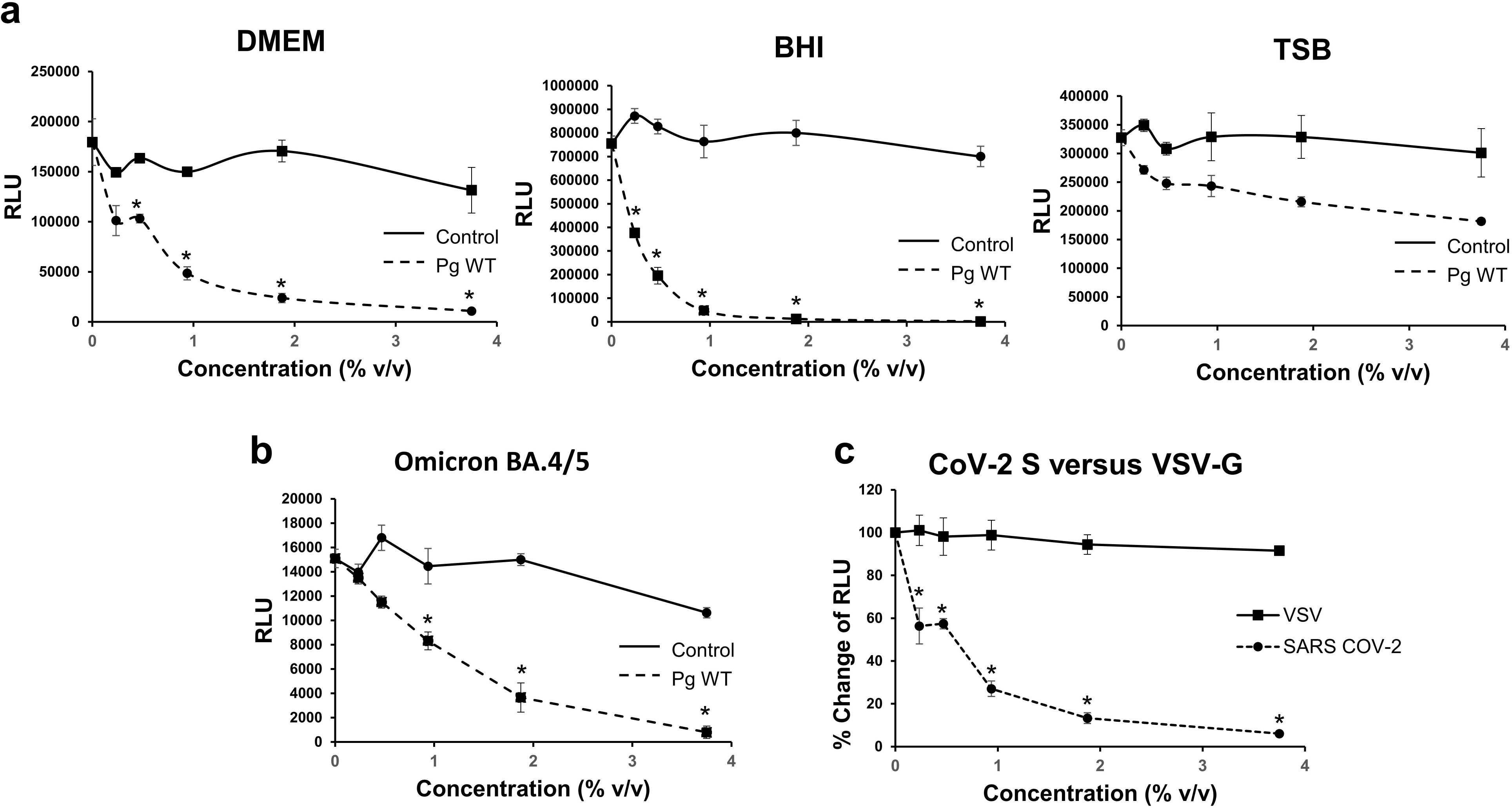
*P. gingivalis* factors inhibit SARS-CoV-2 pseudovirus infection. ACE2-overexpressing 293T cells were infected with single-round HIV-1 lentiviral vector pseudotyped with SARS-CoV-2 (CoV-2) spike (Wuhan strain) or VSV-G. The viruses contain luciferase reporter to allow for measurement of infection in RLU. a. Inhibition of infection by various concentrations (% v/v) of supernatants of *P. gingivalis* W83 WT (*Pg* WT) grown in Brain-Heart Infusion (BHI), Tryptic Soy Broth (TSB) and 10% FCS DMEM medium and medium control (Control) is shown. b. The inhibitory effect of supernatant of *P. gingivalis* grown in 10% FCS DMEM on CoV-2 Omicron BA.4/5 pseudotyped lentivirus was measured. c. Comparison of CoV-2 pseudovirus versus VSV control inhibition by *P. gingivalis* supernatant (grown in DMEM/10%FCS) is shown and inhibition of infection (%Change of RLU) is indicated. The mean luciferase activity ± sd (N=3) of infected cells was determined. Difference between infection with treatment and control as well as between infection with CoV-2 and VSV pseudovirus was considered significant (*P < .05) using Student’s t-test.

Host airway epithelial cells from the lungs are the primary target cells for SARS-CoV-2 infection. Furthermore, periodontal pathogens found in the lungs of patients with certain respiratory conditions could play a role in lung cell infection by SARS-CoV-2.^18^ We, therefore, tested the effects of *P. gingivalis* factors on infection of lung cells, using lung tumor epithelial cell line H838 overexpressing ACE2 as target cells. We found that *P. gingivalis* supernatants at 2% and 4% concentration of the total media volume inhibited infection by roughly 50% (Fig. 2). Altogether, these viral inhibition studies strongly suggest that *P. gingivalis* secretes factors capable of inhibiting SARS-CoV-2 infection that was mediated by the spike protein.

**Fig. 2.**
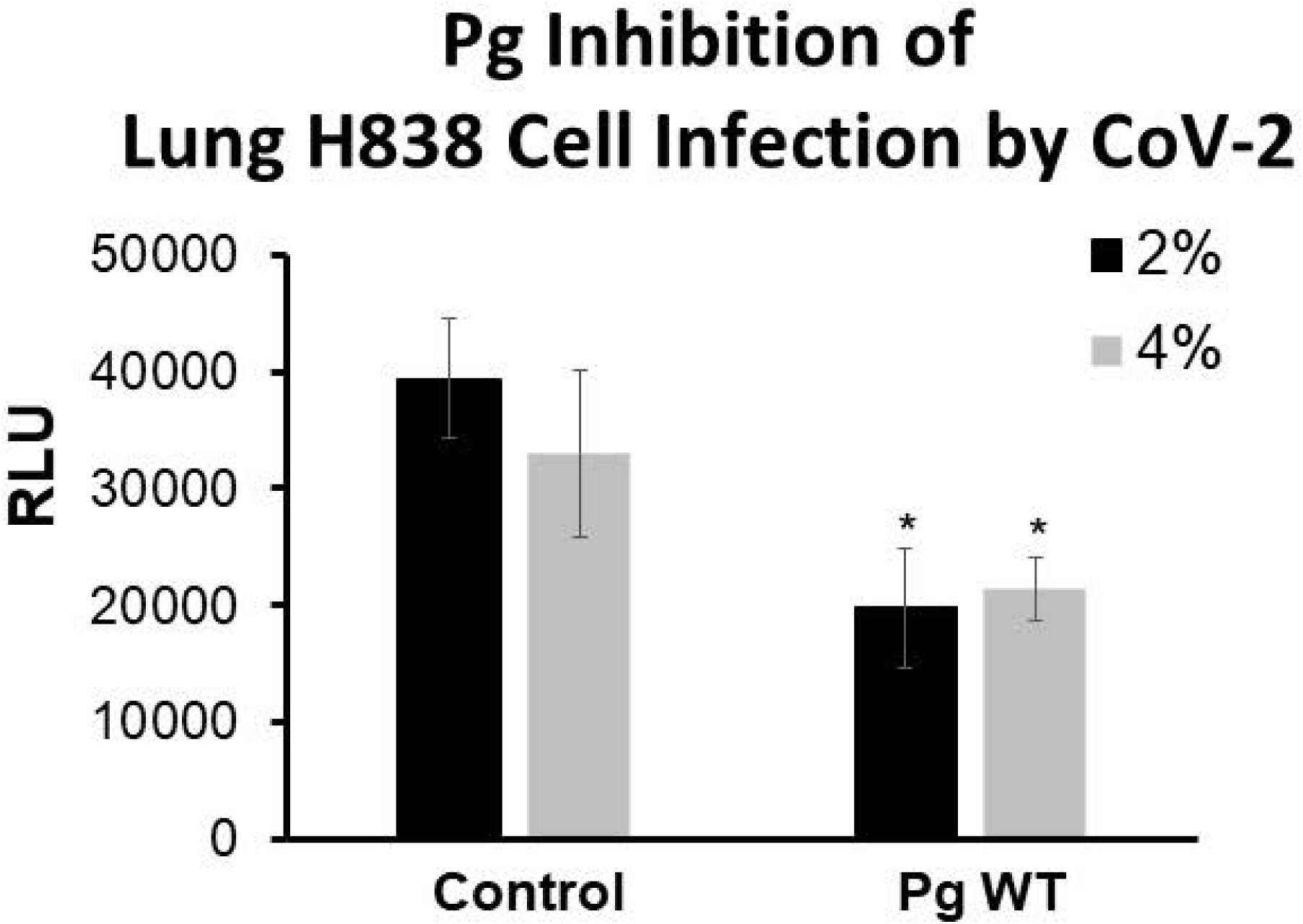
*P. gingivalis* factors inhibit lung cancer cells infection by SARS-CoV-2 pseudovirus. The lung cancer cell line H838 was transiently transfected to overexpress ACE2. Supernatants of *P. gingivalis* wt grown in DMEM/10% FBS at 2% and 4% v/v were able to inhibit infection by SARS-CoV-2 pseudovirus transducing luciferase. Infection was measured by luciferase activity (RLU) and the mean RLU ± sd (N=3) of infected cells with or without *P. gingivalis* supernatant was determined. Difference between infection with treatment and control was considered significant (*P < .05) using Student’s t-test.

### *P. gingivalis* PGDHC inhibited SARS-CoV-2 pseudovirus infection

To investigate which secreted factors were inhibiting SARS-CoV-2 pseudovirus infection, we tested known compounds that have been shown to be immunomdulatory including phosphoglycerol dihydroceramides (PGDHC) and LPS. We found that PGDHC strongly inhibited SARS-CoV-2 pseudoviruses containing the spike protein (IC50 ∼2 ug/ml) (Fig. 3a), while the LPS had no effect on the pseudoviral infection (Fig. 3a). PGDHC did not inhibit VSV pseudoviruses, which suggests that PGDHC inhibition of SARS-CoV-2 was mediated by the spike protein (Fig. 3b). Interestingly, supernatants from the SPT-mutant, ^27^ which does not produce PGDHCs or phosphoethanoloamine dihydroceramides (PEDHCs) had similar inhibitory activity as the parent strain (Fig. 1a and Fig. 3c). Taken together, these results suggest that PGDHC and other unknown factors produced by *P. gingivalis* are strong inhibitors of SARS-CoV-2 infection that is mediated by the spike protein.

**Fig. 3.**
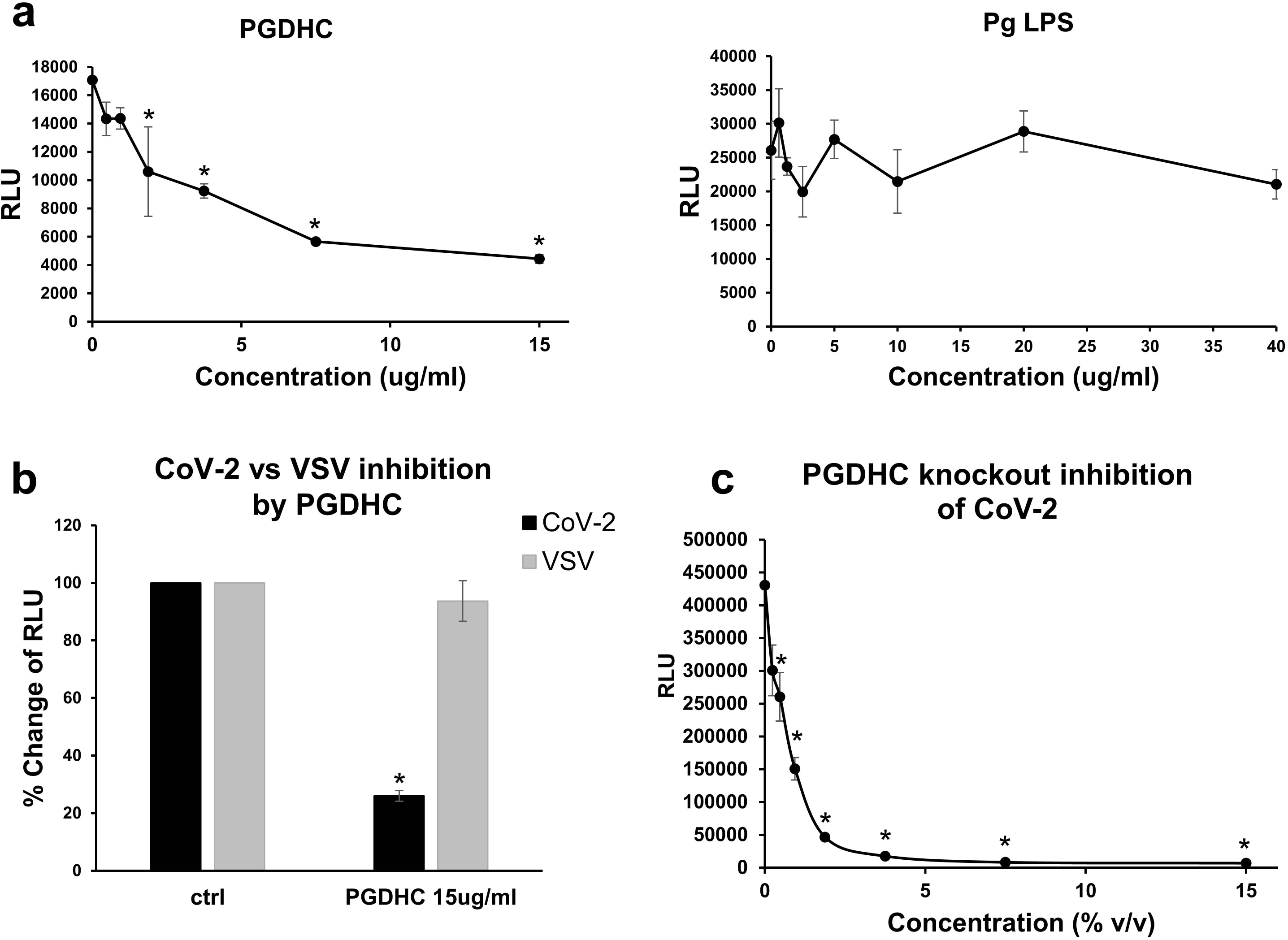
*P. gingivalis* phosphoglycerol dihydroceramide (PGDHC) inhibits SARS-CoV-2 pseudovirus infection. a. ACE2+ 293T cells were pre-incubated for 6 hours with various concentrations of *P. gingivalis* PGDHC or LPS and subsequent infection in the presence of these *P. gingivalis* factors with SARS-CoV-2 pseudovirus was assessed. b. Inhibition by PGDHC on infection (indicated as % Change of RLU) was specific to SARS-CoV-2 (CoV-2) but not VSV-pseudovirus (VSV). Infected ACE2+ 293T cells with PGDHC (15ug/ml) or without (ctrl) are shown. c. Effects of PGDHC gene knockout on SARS-CoV-2 pseudoviral inhibition. Infected ACE2+ 293T cells in the presence of various concentrations of PGDHC knockout supernatant are shown. Infection was measured by luciferase activity (RLU) and the mean RLU ± sd (N=3) of infected cells with or without *P. gingivalis* supernatant was determined. Difference between infection with treatment and control as well as between CoV-2 and VSV pseudovirus infection was considered significant (*P < .05) using Student’s t-test.

### *P. gingivalis* gingipains inhibited SARS-CoV-2 pseudovirus infection

Gingipains, which are a family of cysteine proteases secreted by *P. gingivalis*, have previously been shown to have antiviral effects towards HIV^25^ and have been associated with the increased metapneumovirus infection.^28^ To assess the possible role of gingipains in COVID-19 infection, we used the triple gingipain knockout *P. gingivalis* strain (*RGPA, RGPB* and *KGP* knockout). We found that the antiviral effects of *P. gingivalis* wildtype supernatants were dramatically reduced in the *P. gingivalis* gingipain knockout supernatants (Fig. 4a and b). In addition, the triple gingipain knockout supernatant at higher concentrations (7.5% and 15% v/v) still significantly inhibited SARS-CoV-2 pseudoviral infection of ACE2+ 293T cells (Fig. 4b) despite the loss of cytotoxic effects at these high supernatant concentrations (Fig. 4c). Taken together, these results suggest that gingipains mediate the inhibition of SARS-CoV-2 pseudovirus through its cytotoxic effects. Also, in addition to gingipains, there are other possible anti-SARS-CoV-2 mediators present in *P. gingivalis*.

**Fig. 4.**
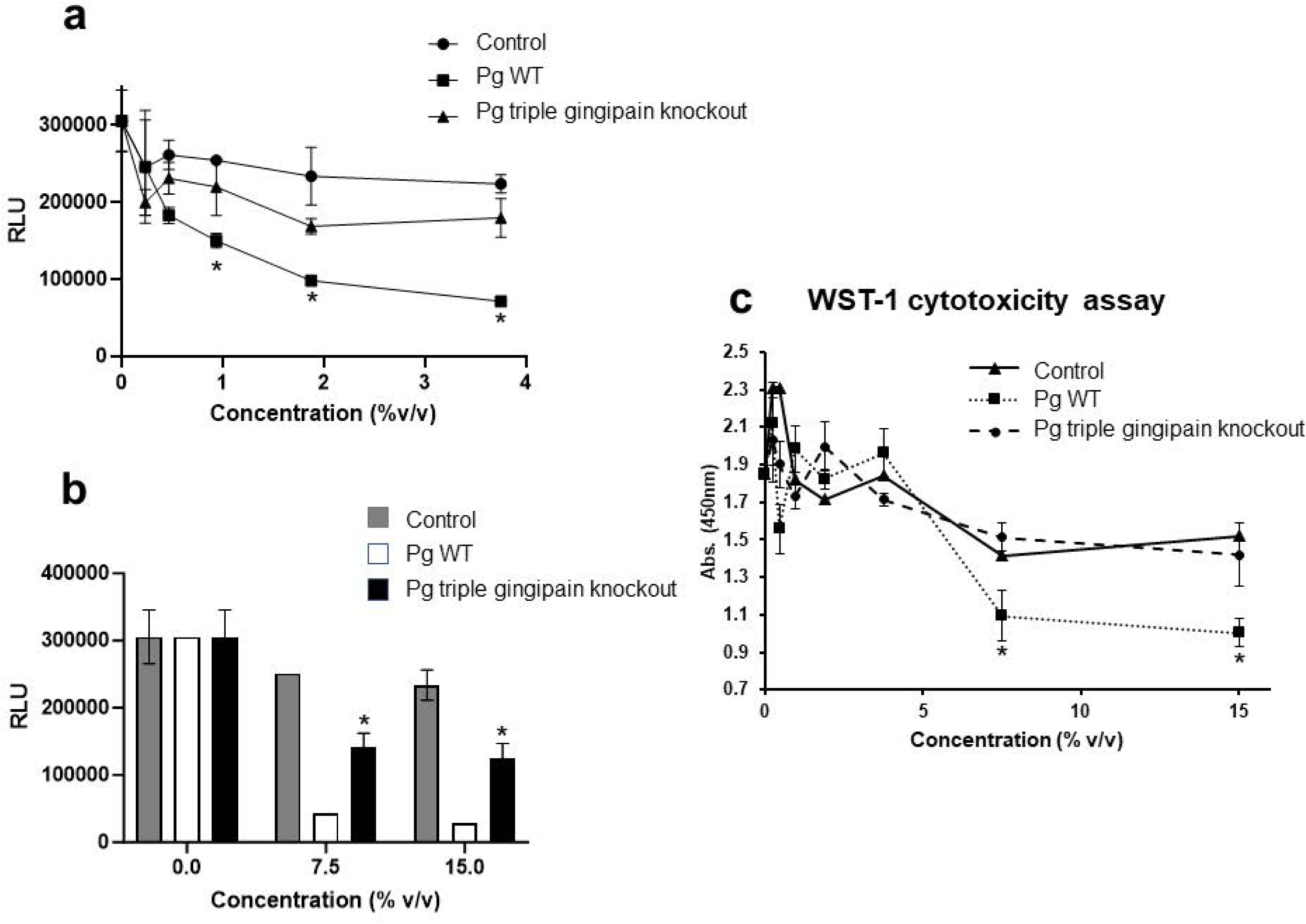
Effects *P. gingivalis* gingipains in SARS-CoV-2 pseudovirus infection. a. For SARS-CoV-2 pseudovirus inhibition of ACE2+ 293T infection, various concentrations (%v/v) of supernatants from wildtype *P. gingivalis* (*Pg* WT) and triple gingipain knockout *P. gingivalis* mutant (*Pg* W83^Δ^*^kgp^*^Δ^*^rgpA^*^Δ^*^rgpB^*), and 10% FCS DMEM only control (DMEM ctrl) was used. b. Inhibition of ACE2+ 293T infection at higher bacterial supernatant concentrations (7.5% and 15%) is shown. c. Cytotoxicity of ACE2+ 293T (measured by WST) at high concentrations of *Pg* WT supernatant but not triple gingipain knockout is shown. Infection was measured by luciferase activity (RLU) and the mean RLU ± sd (N=3) of infected cells with or without *P. gingivalis* supernatant was determined. Difference between infection with treatment and control was considered significant (*P < 0.05) using Student’s t-test.

### *P. gingivalis* antiviral factors inhibited SARS-CoV-2 infection independent of spike protein binding to ACE2 receptor

We evaluated whether the inhibition of SARS-CoV-2 pseudoviral infection occurred during the early steps of the viral life cycle. To assess the effects on viral entry, we determined the ability of *P. gingivalis* supernatant and purified PGDHCs to interfere with the binding of SARS-CoV-2 spike protein to ACE2 receptor protein using Biolayer Interferometry analysis. As expected, the spike protein bound to ACE2 efficiently (Fig. 5a). Neither the *P. gingivalis* supernatant (15% v/v) nor PGDHCs (15ug/ml) affected the binding of the purified spike protein to purified ACE2 receptor protein (Fig. 5a).

**Fig. 5.**
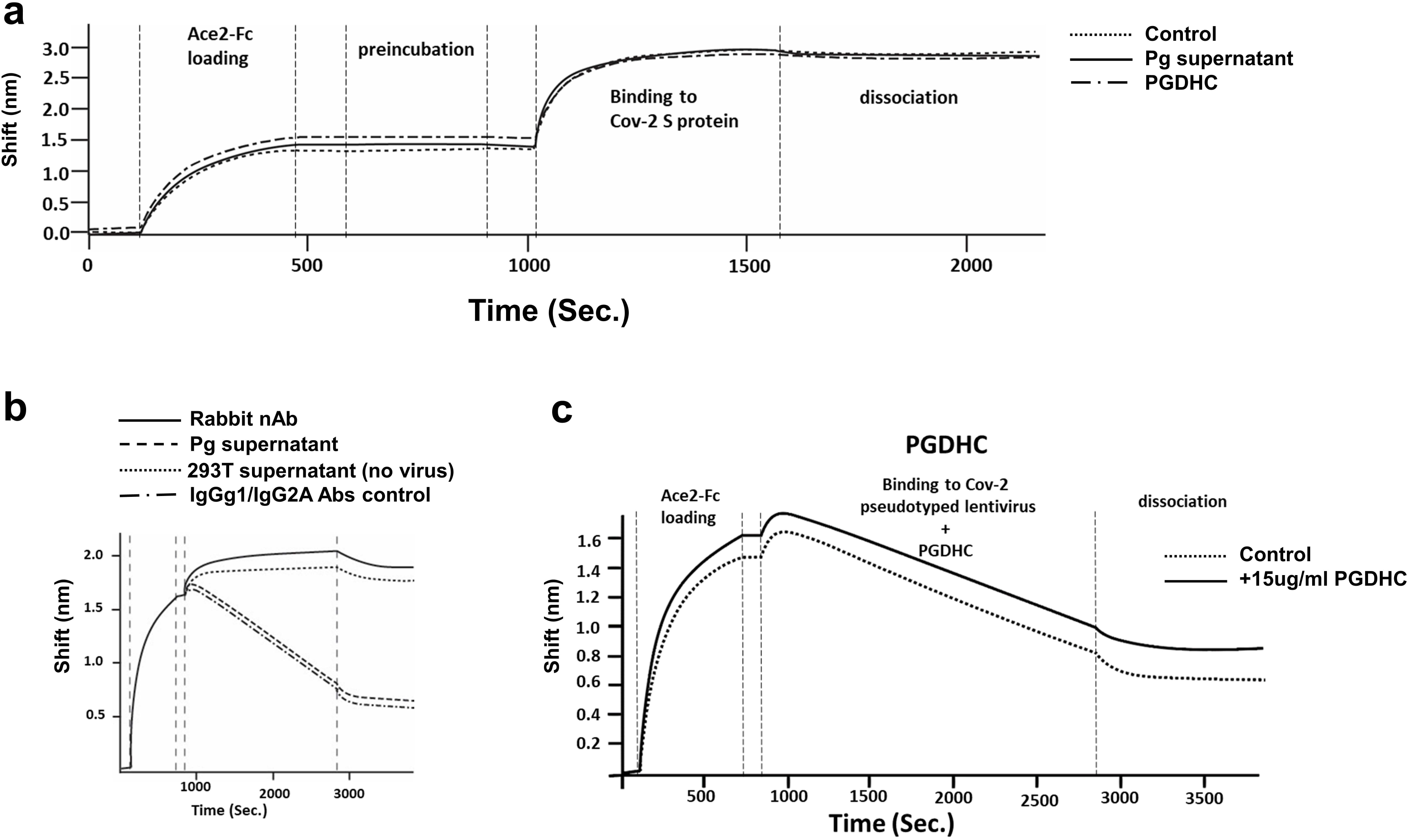
*P. gingivalis* PGDHC and factors do not affect binding of SARS-CoV2 S protein to ACE2. a. BLI binding assay was performed to determine if PGDHCs, *P. gingivalis* supernatant or medium alone (Control) inhibited binding of S protein to ACE2-Fc. b. SARS-CoV-2 pseudovirions binding to the ACE2 loaded biosensor treated with Rabbit neutralizing antibody (Rabbit nAb), *P. gingivalis* supernatant, non-neutralizing IgGg1/IgG2A Abs control or 293T supernatant only (no virus). c. SARS-CoV-2 pseudovirions binding to the ACE2 loaded biosensor treated with 15ug/ml PGDHC or without PGDHC (Control). The BLI curves of the binding as well as the loading, preincubation, binding and dissociation steps are shown. The sensogram shows the binding shift (nm) over time (seconds).

Using BLI analysis, we also assessed the ability of *P. gingivalis* secreted factors to inhibit the binding of pseudoviruses containing the spike protein to purified ACE2 protein. Binding of the pseudovirus to ACE2 caused a negative shifting of the BLI curve similar to prior observations with virus binding (Fig. 5b).^29,30^ Using an anti-spike rabbit neutralizing antibody (nAb), the binding of pseudovirus containing the spike protein to ACE2 receptor was inhibited based on the lack of negative shifting of the BLI curve (Fig. 5b). By contrast, the *P. gingivalis* supernatant, PGDHC and non-neutralizing antibodies did not inhibit SARS-CoV-2 pseudovirus binding to ACE2 (Fig. 5b and c). Thus, these results altogether suggest that the observed inhibition by *P. gingivalis* is not mediated by the binding of the spike protein on the virus to ACE2 receptor.

### *P. gingivalis* inhibited SARS-CoV-2 infection at the fusion step of the virus life cycle

We assessed whether the secreted factors produced by *P. gingivalis* interfered with the virus life cycle, specifically after the binding of spike protein to ACE2. A syncytium formation assay we developed was used to assess whether *P. gingivalis* is inhibiting SARS-CoV-2 pseudoviral infection at the viral-host cell fusion step of the virus life cycle. To develop the syncytium-formation assay, we generated TZM.bl expressing SARS-CoV-2 S protein and 293T cells expressing a Tat-P2A-Rev (Fig. 6a, left panel). Once co-cultured, the spike/ACE2 interaction resulted in membrane fusion and the formation of syncytium (Fig. 6a, left and middle panels). In the fused cells, Tat transactivates the LTR-Luc cassette present in the TZM.bl cells which results in the expression of the luciferase reporter gene (Fig. 6a, left and right panels). As expected, addition of anti-spike protein mouse neutralizing antibody inhibited syncytium formation at an IC50 of 0.5ug/ml (Fig. 6a, right panel).

**Fig. 6.**
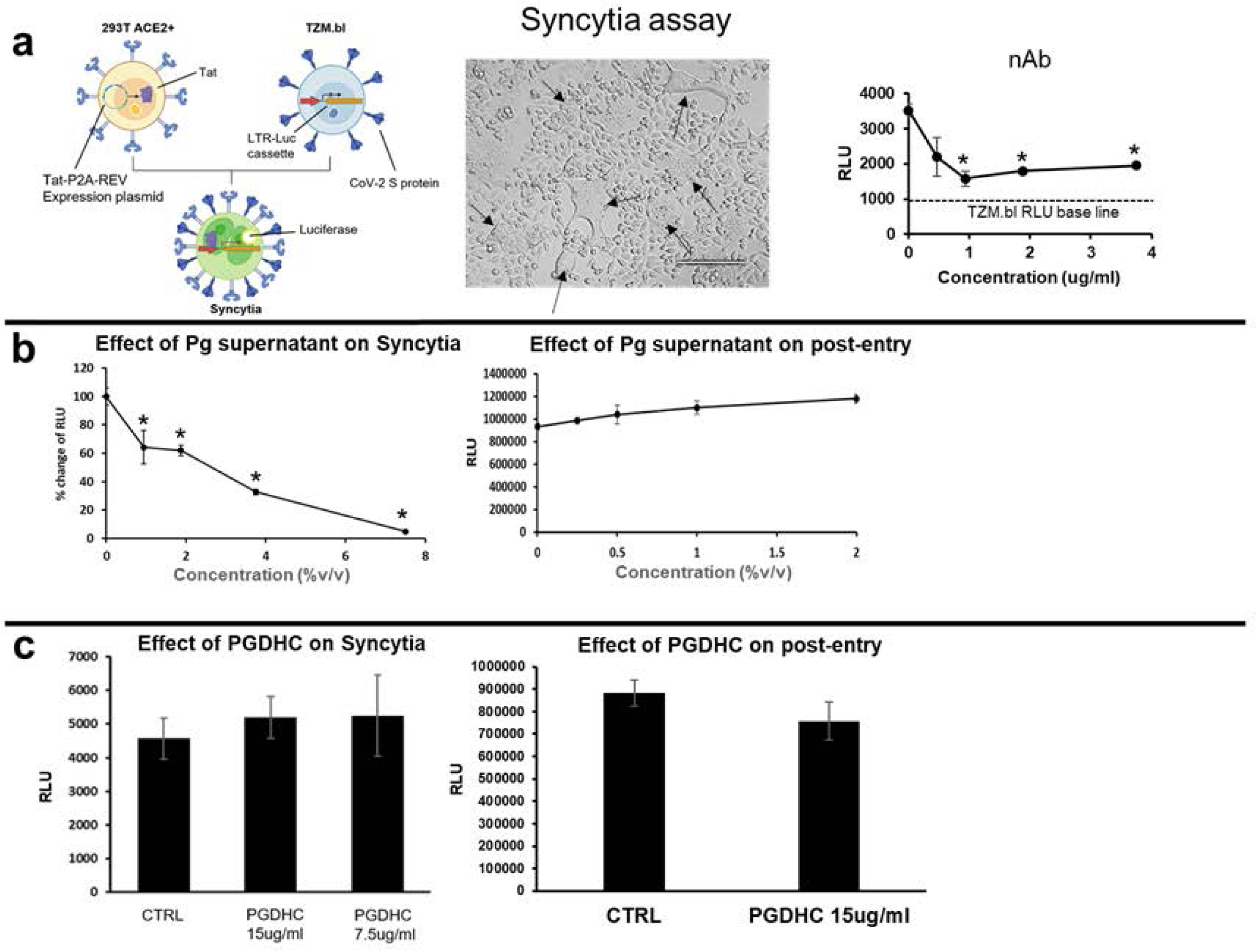
Effect of *P. gingivalis* factors on fusion and postentry steps of the viral life cycle. A syncytium formation assay was used to assess the effects of *P. gingivalis* factors on the fusion step of the viral life cycle. a. Luc activity (RLU) indicates fusion between 293T ACE2+ and Tzm-bl cells expressing the S protein (left panel). Light microscopy of syncytia formed (indicated by arrows) is shown (middle panel). Inhibition of syncytium formation by rabbit anti-S neutralizing antibody (nAb) is shown (right panel). b and c. The effects of *P. gingivalis* supernatant at various concentrations (%v/v) and PGDHC at 7.5 ug/ml or 15 ug/ml on syncytium formation (left panels) and postentry (right panels) are shown. 10% FCS DMEM only was used as control (CTRL)To measure the effects of *P. gingivalis* factors on postentry, ACE2+ 293T were infected and 24 hours later the *P. gingivalis* factors were added and RLU was measured 2 days later. Infection was measured by luciferase activity (RLU) and the mean RLU ± SD (N=3) of infected cells was determined. Difference between infection with treatment and control was considered significant (*P < .05) using Student’s t-test.

We observed that *P. gingivalis* supernatant but not PGDHCs was capable of inhibiting the formation of syncytium (Fig. 6b and c, respectively). *P. gingivalis* supernatant potently inhibited syncytium formation with an IC50 ∼2.7% v/v. A post-entry assay showed that neither the *P. gingivalis* supernatant nor PGDHC had an effect on pseudoviral infection at the post-entry step (Fig. 6b and c, respectively). Altogether, these results strongly suggest that *P. gingivalis* anti-COVID-19 factors but not PGDHC inhibit SARS-CoV-2 infection mediated by the spike protein at the fusion step of the viral life cycle. Additionally, the secreted factors had no effect on viral post-entry process, specifically once the virus has already entered the host cell.

### SARS-CoV-2 pseudovirus infection is inhibited by *P. gingivalis* and other oral bacteria

Because the oral pathogen *P. gingivalis* showed some anti-SARS-CoV-2 properties, it is possible that other oral bacteria, either pathogens or commensals would exhibit similar properties. To explore this possibility, oral bacterial supernatants from different species belonging to various microbial complexes classified by Dr. Socransky were generated and tested for anti-SARS-CoV-2 properties.^31^ We observed that supernatants from oral pathogens *Actinobacillus* a*ctinomycetemcomitans* (Y4 strain) and *Actinobacillus* a*ctinomycetemcomitans* (29523 strain) as well as the oral commensal *Actinomyces odontolyticus* inhibited pseudoviral infection.^32^ Strong inhibition by these oral bactera was observed at very low supernatant concentrations (Fig. 7). Cytotoxicity analysis using WST-1 assay indicated that these oral pathogens inhibited SARS-CoV-2 pseudoviral infection due to the highly cytotoxic effects of the supernatants on ACE2 293T target cells. In contrast, supernatants from cultures of other oral commensals including *S. mitis* and *S. gordonii* had poor antiviral effect (Fig. 7). These results suggest that SARS-CoV-2 infection can be inhibited by certain oral bacteria including *P. gingivalis, A. actinomycetemcomitans, and A. odontilyticus*; yet not all oral microbiota exhibit this inhibitory activity.

**Fig. 7.**
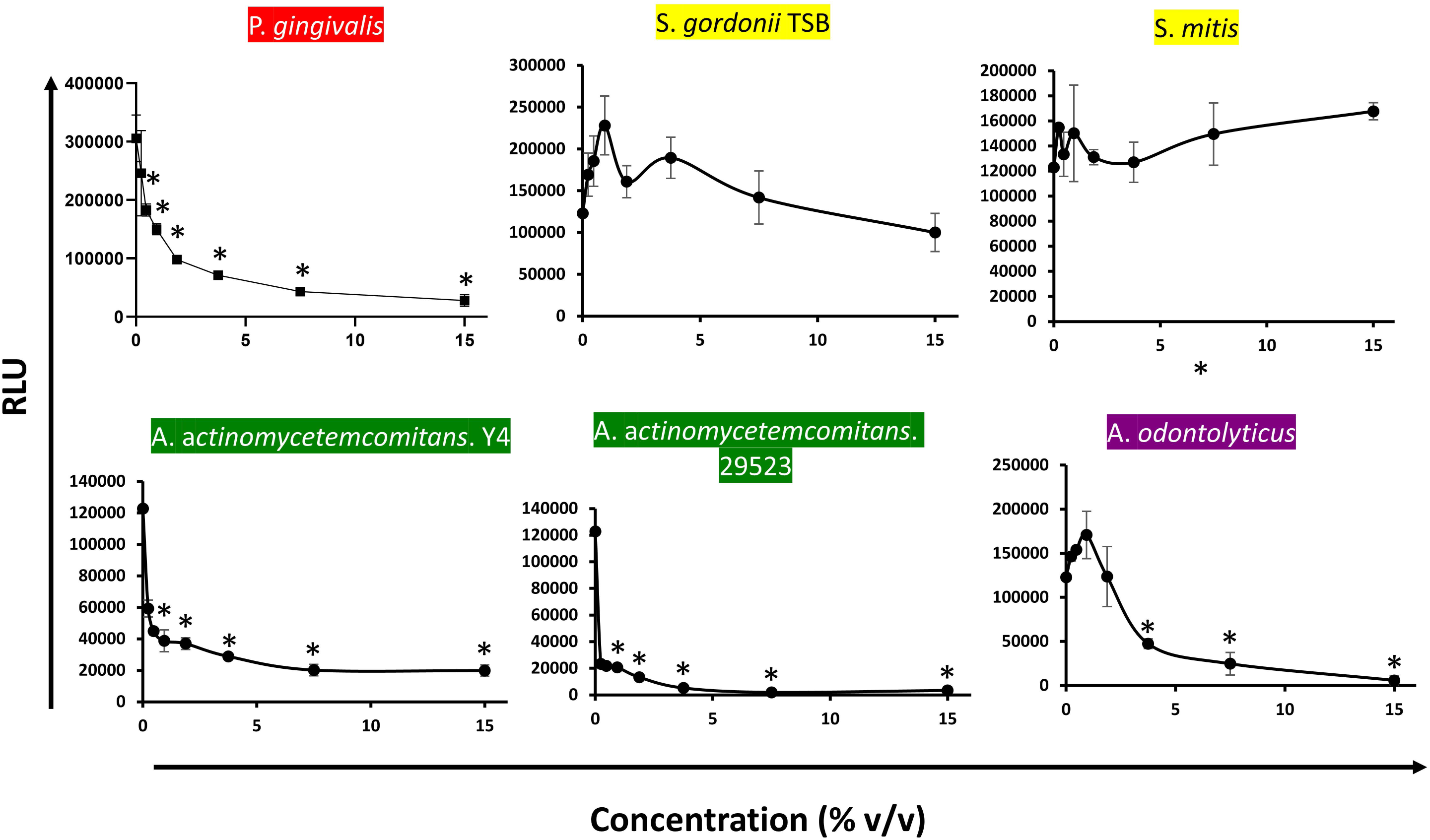
Oral bacteria belonging to various microbial complexes can inhibit SARS-CoV-2 pseudoviruses. The red complex (*P. gingivalis*), yellow (*S. gordoni*and *S. mitis*), green (*A. actinomycetemcomitans Y4 and A. actinomycetemcomitans 29523*) and purple (*A. odontilyticus*) grown in 10% FCS DMEM except for *S. gordonii* (grown in TSB) were tested for modulatory activity against SARS-CoV-2 pseudovirus infection. Oral bacterial supernatants at various concentrations were added to SARS-CoV-2 pseudovirus infection assay using ACE2+ 293T target cells. Infection was measured by luciferase activity (RLU) and the mean RLU ± SD (N=3) of infected cells was determined. Difference between infection with treatment and control was considered significant (*P < .05) using Student’s t-test.

### Oral bacteria from healthy individuals modulated SARS-CoV-2 infection

Since we observed that secreted factors from certain type of strains propagated in the laboratory had anti-SARS-CoV-2 effect, we explored whether oral bacteria enriched from saliva samples from healthy human volunteers exhibited any modulatory activity towards SARS-CoV-2. Interestingly, out of all cultures grown from 37 individuals, those from 9 individuals exhibited antiviral activity, while culture supernatants from 17 individuals enhanced infection (Fig. 8). Moreover, 11 out of the 37 healthy volunteers harbored oral bacteria with no effect on pseudovirus infection (Fig. 8). Also, the anti-SARS-CoV-2 property of bacterial supernatants from three individuals was associated with strong cytotoxicity to target cells (data not shown). Collectively, these results strongly suggest that oral bacteria present in healthy human individuals have the capacity to modulate SARS-CoV-2 pseudoviral infection. Specifically, the modulatory activity of oral bacteria present in the saliva can either be inhibitory, enhancing or neutral to SARS-CoV-2 infection.

**Fig. 8.**
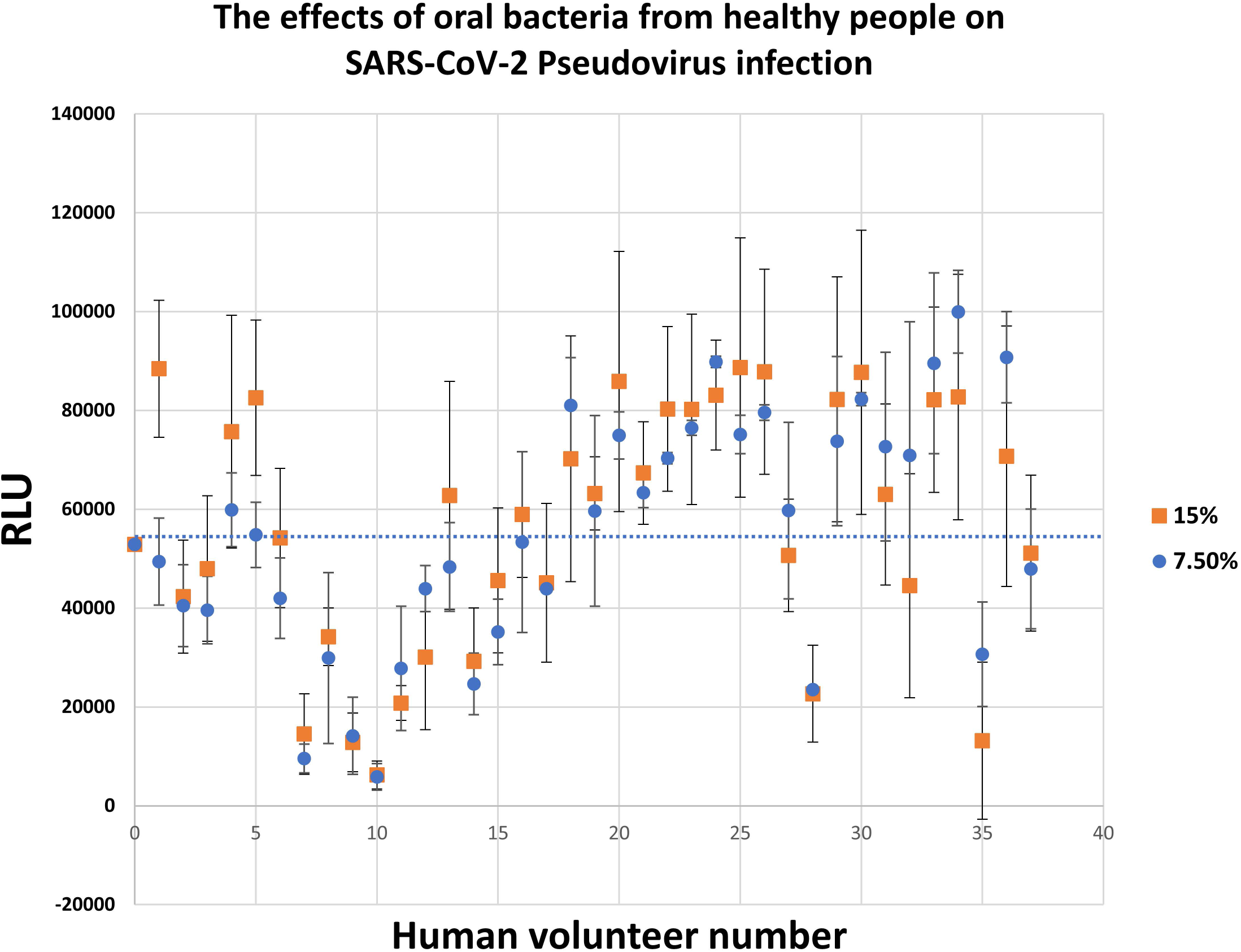
Oral bacteria from healthy people modulate SARS-CoV-2 pseudovirus Infection. Oral bacteria were isolated from saliva of 37 healthy human volunteers and tested for modulatory activity against SARS-CoV-2 pseudovirus infection. Oral bacteria were cultured aerobically in BHI and supernatants at either 7.5% or 15% v/v concentration were tested for ability to inhibit SARS-CoV-2 pseudovirus infection of ACE2+ 293T cells. Infection was measured by luciferase activity (RLU) with the mean RLU ± SD (N=3) of infected cells. The bacterial supernatant activity towards SARS-CoV-2 from each volunteer is shown. The baseline level of infection is indicated by the dotted line at approximately 55,000 RLU.

## Discussion

The role the oral microbiome plays in COVID-19 infection has not been elucidated, and in this study, we demonstrated a potential influence of the oral microbiome on SARS-CoV-2 infection. Interestingly, using an in vitro pseudovirus infection assay, we demonstrated that certain oral bacteria, specifically two oral pathogens (*P. gingivalis* and *A. actinomycetemcomitans)* and a commensal *(A. odontilyticus)* were capable of inhibiting SARS-CoV-2. In contrast, two oral commensals that were tested, specifically *S. mitis* and *S. gordonii* had no detectible antiviral properties. Interestingly, supernatants from enrichment cultures generated with saliva samples from healthy human individuals were either capable of inhibiting or enhancing while some had no effect on SARS-CoV-2 pseudovirus infection.

The anti-SARS-CoV-2 effect of Pg was noteworthy in that we observed a marked antiviral effect exerted by *P. gingivalis* supernatant on SARS-CoV-2 pseudoviral infection of ACE2+ 293 T cells with an IC50 of 0.3% v/v. *P. gingivalis* was not only capable of inhibiting pseudoviruses expressing the spike protein from the original Wuhan strain but also the BA.4/5 omicron variant. In addition, we found that *P. gingivalis* also inhibited the infection of relevant lung cells (adenocarcinoma cell line H838), which is important since airway lung epithelial cells are the in vivo target cells of SARS-CoV-2. Periodontal pathogen infections of the lungs have been observed and it is proposed that translocation of periodontal pathogens from the oral cavity into into the lungs could influence SARS-CoV-2 infection of lung epithelial cells.^18,19^ If *P. gingivalis* were to be found in the lungs of patients with COVID-19, *P. gingivalis* could potentially inhibit SARS-CoV-2 infection of lung epithelial cells.

Inhibition of ACE2+ H838 was significant but not as potent as the 293T cell target cells, which suggests that there are differences in the expression of cellular factors in lung versus 293T cells that are important for infection. The use of primary lung cultures/organoids or lung infection in animal models is needed to assess fully the antiviral property of *P. gingivalis*. In addition to cellular factors expressed on target cells, expression of bacterial factors with antiviral properties will be key. We found that bacteria grown in DMEM/10%BSF and BHI but not TSB are capable of inhibiting SARS-CoV-2 pseudovirus, suggesting that perhaps, the growth conditions in the oral cavity could play are role in the production of antiviral factors by *P. gingivalis*.

We determined that purified PGDHCs from *P. gingivalis* strongly inhibited SARS-CoV-2 pseudovirus infection. In contrast, *P. gingivalis* LPS, known to activate several cells of the immune system interacting with TLR4 had no anti-SARS-CoV-2 property.^33^ To provide context, the different subtypes of *P. gingivalis* dihydroceramides (PGDHCs and PEDHCs) have been shown to be secreted into the environment on outer membrane vesicles (OMVs) and be immunomodulatory.^34,35^ In addition, these dihdyroceramides are thought to be virulence factors that are associated with bone loss in periodontitis patients as well as with the pathogenesis of Alzheimer’s.^36-38^ Although the underlying mechanisms involved in the PGDHC-mediated viral inhibition still need to be determined, it is possible that the purified PGDHCs because of their sphingolipid structure are inhibiting the virus membrane-to-cell membrane fusion or the virus decapsulation by modifying the lipid bilayer biophysical properties, by acting on the ACE2-viral membrane glycoprotein complex or acting on accessory proteins required for the successful virus entry. PGDHCs have also been shown to trigger signal transduction pathways involved in osteoclastogenesis and amyloidogeneis, which may inhibit SARS-CoV-2 infection.^36-38^

Yet, it is important to note that like the wild-type, the SPT mutant was also strongly inhibitory, which is contradictory, suggesting that synthesis of PGDHCs by *P. gingivalis* is not required for viral inhibition. However, while it is true that this mutant does not synthesize PGDHCs, the SPT mutant is also defective in the synthesis of other subtypes of DHCs, including the phosphoethanolamine dihydroceramides (PEDHCs). Therefore, it is possible that the different DHC types may promote or constrain invasion and act so by different mechanisms. Lastly, it may be that in the purified form, PGDHCs are inhibitory but they are not inhibitory when delivered as an integral part of an OMV membrane. Hence, further studies are required to elucidate the mechanisms involved in the observed PGDHC-mediated viral inhibition.

*P. gingivalis* gingipains also inhibited SARS-CoV-2 pseudovirus infection based on our observation that the triple gingipain knockout *P. gingivalis* strain had a significantly reduced ability to inhibit infection. Since we did not test purified gingipains, it is possible that other proteins regulated by gingipain aminopeptidase activity could be responsible for inhibiting SARS-CoV-2 pseudovirus infection.^39^ Gingipains were previously shown to be critical in processing fimbriae and several bacterial proteins that contribute to hemagglutination, coaggregation, and hemoglobin binding (reviewed in reference ^40^). Gingipain mediated inhibition is likely due to its known host cell cytotoxicity property,^41,42^ consistent with our observation. RGPA and KGP were shown to inhibit HIV by binding the HGP44 domain to the HIV envelope glycoprotein^25^ and the ability of catalytically active gingipains to bind to the spike protein to inhibit infection should be explored.

In addition to the known compounds such as PGDHC and gingipains, there are other unidentified anti-SARS-CoV-2 factors produced by *P. gingivalis* based on our observation that PGDHC and triple gingipain knockout mutants retained significant antiviral activity. Our entry and post-entry assays revealed that these unknown factors present in *P. gingivalis* supernatants inhibited SARS-CoV-2 pseudovirus at the fusion step of the virus life cycle. Inhibition of fusion was likely through the spike protein since the supernatant had no effect on infection of 293T ACE2+ cells by control VSV pseudovirus. While fusion was inhibited, entry through binding of spike protein or pseudovirus to ACE2 was not inhibited by the supernatants nor by purified PGDHCs. The antiviral factors present in *P. gingivalis* supernatant could interfere with fusion of the viral membrane with the cellular membrane following S trimer-ACE2 interaction but by what mechanism needs to be determined.

The potential for oral bacteria to modulate SARS-CoV-2 infection is entirely possible in that bacteria are known to possess antiviral compounds that have been shown disrupt the viral life cycle by interfering with viral attachment and penetration, genome integration, replication and transcription.^20,21^ The ability of *P. gingivalis* supernatants to inhibit SARS-CoV-2 suggests that SARS-CoV-2 inhibitor(s) are likely soluble secreted factors that can inhibit the fusion step of the SARS-CoV-2 life cycle. Metabolomic, proteomic, and biochemical studies are currently underway to identify the antiviral compounds produced by *P. gingivalis*.

A previous study showing the association between periodontitis and COVID-19 disease severity suggests that the periodontal pathogens including *P. gingivalis* may enhance infection or exacerbate disease.^43-45^ This observation seems inconsistent with our present findings showing that *P. gingivalis* could inhibit SARS-CoV-2 infection. One plausible explanation to reconcile these two seemingly contradictory findings is that the association between periodontitis and severe disease may not be due to the direct effect of *P. gingivalis* on SARS-CoV-2 infection, but rather, due to the inflammation associated with periodontitis. Inflammation elicited by *P. gingivalis* has been showed to trigger apoptosis in H1N1 infected lung cells.^46^ Thus, *P. gingivalis* could directly inhibit SARS-CoV-2 infection per our observation, but it could also promote severe disease by its ability to induce inflammation in people with COVID-19. Indeed, excessive inflammation seen in patients experiencing a “cytokine storm” is strongly associated with severe disease.^43,47,48^

In addition to *P. gingivalis*, other oral bacteria could play a role in modulating SARS-CoV-2 infection. We tested supernatants isolated from oral bacteria clones representing the various microbial complexes determined by Socransky and coworkers that classify oral commensals and pathogens.^31^ We found that the oral pathogen A. a*ctinomycetemcomitans* and the commensal bacterium A. *odontolyticus* inhibited pseudoviral infection mainly due to their highly cytotoxic effects on ACE2+ 293T target cells. By contrast, oral streptococci such as *S. mitis* and *S. gordonii* had no antiviral effect. It is possible that oral microbiota could also exert their antiviral effects indirectly through their ability to alter gene expression and hence the susceptibility of target cells.^48,49^

Lastly, we explored the possible role of the oral microbiome in COVID-19 by assessing the modulatory effect of supernatants generated from microbial communities directly enriched from saliva of healthy human volunteers. Surprisingly, we found that oral bacteria present in healthy human individuals have differing capacity to modulate SARS-CoV-2 pseudoviral infection. Individuals harbor oral bacteria that were either inhibitory, enhancing or inactive to SARS-CoV-2 infection. We are currently investigating what explains this phenomenon. It could be due to differences in oral microbiome composition and host-microbiome interactions that could regulate the production of these virus modulatory compounds.^49^

Given the importance of the oral microbiome in health and disease, we began to address the role of the oral microbiome in COVID-19, a disease that clearly remains a global public health problem.^50^ Our in vitro studies revealed striking differences between oral bacteria species in that certain periodontal pathogens, especially *P. gingivalis* produce known and unknown compounds that are capable of inhibiting SARS-CoV-2. Another conclusion from this study is that differences in the oral microbiome composition could explain why certain people may have modulating activities. It can be proposed that these inhibiting or enhancing activities could contribute significantly to either the resistance or susceptibility of people to SARS-CoV-2 infection. More studies are needed to explore these interesting questions regarding the role of the oral microbiome in COVID-19. Ultimately, a clear understanding of how the oral microbiome plays a role in COVID-19 will lead to an effective therapy to treat or prevent infection.

## Materials and Methods

### Saliva Collection

De-identified unstimulated saliva was collected from 37 healthy human volunteers, ages 18 and older. All participants understood the nature of the study and provided informed consent. The Nova Southeastern University Institutional Review Board approved the study design as well as the procedure for obtaining informed consent (NSU IRB protocol number 2020-607). All experiments were performed in accordance with the approved guidelines.

### Mammalian cell cultures

293T (ATCC CLR-3216), ACE2+ 293, H838 (ATCC CLR-5844), and TZM.bl cells were cultured in DMEM complemented with 10% SBF and 2% Penicillin (5000 U/ml)/Streptomycin (5mg/ml) (GIBCO, Waltham, MA, USA), and maintained at 37°C in 5% CO2 atmosphere. The TZM.bl cells were obtained from the NIH HIV Reagent Program, Division of AIDS, NIAID, NIH. TZM-bl Cells, ARP-8129, were contributed by Dr. John C. Kappes, Dr. Xiaoyun Wu and Tranzyme Inc.

### Bacterial cultures

*Porphyromonas gingivalis* (*Pg* WT) W83 BAA-308 was purchased from American Type Culture Collection (Manassas, VA, USA). *P. gingivalis* RGPA/RGPB/KGP triple knockout bacteria were provided by Jan S. Potempa, University of Louisville, KY, USA. A serine palmitoyl transferase (SPT) knockout strain of *P. gingivalis* W83 which does not produce any dihydroceramides (sphingolipid-null strain) was provided by Mary Ellen Davey, The ADA Forsyth Institute, Cambridge, MA, USA. Bacteria were cultured on a sheep blood agar plate at 37 °C under anaerobic condition using the Anaeropack system (Mitsubishi Gas Chemical, Tokyo, Japan). Once colonies were detected, the bacteria were transferred in 10ml liquid growing medium in the same anaerobic conditions. To grow bacteria from saliva, cell debris was first removed by brief centrifugation and subsequently, 10ul of saliva was cultured in 10ml liquid medium. After 24 hours, bacteria supernatants were collected, filtered supernatant and stored at - 80°C.

### Isolation of PGDHC from *P. gingivalis*

Phosphoglycerol dihydroceramide (PGDHC) lipid class from *P. gingivalis* (ATCC #33277, type strain) was isolated as previously described.^51^ Briefly, *P. gingivalis* was grown in basal (peptone, trypticase and yeast extract, (BBL)) medium supplemented with hemin and menadione (Sigma-Aldrich, St. Louis, MO, USA) and brain heart infusion. Total lipids were extracted from bacterial pellet samples using the phospholipid extraction procedure of Bligh and Dyer as modified by Garbus.^52,53^ Total lipids were fractionated by a two-step HPLC method as previously described.^51,54^ Total lipids were separated first by semi-preparative HPLC separation using a neutral HPLC solvent (hexane:isopropanol:water, 6:8:0.75, v/v/v) followed by LC-MS evaluation of each fraction (see below), and pooling of fractions enriched for PG DHC lipids. The LC-MS analysis was performed using a Sciex500 UHPLC-qTOF instrument and lipid classes were quantified using the multiple reaction monitoring detection mode for negative ions. The pooled fractions were then re-fractionated using the same HPLC solvent supplemented with 0.1% acetic acid. The fractions containing PGDHC were identified by LC-MS and were pooled to yield a highly enriched PG DHC lipid class as previously described.^54^ This method provided highly enriched lipid fractions (>97% purity as determined by mass spectrometric and NMR analysis),^54^ and the purified lipid fractions were devoid of lipoprotein contaminants.

### Soluble SARS-CoV-2 spike protein production

Plasmid containing the SARS-CoV-2 S ectodomain with mutated Furin cutting site and two prolines to enhance stability that was described previously ^55^ was kindly donated by Dr. Jason S. McLellan from The University of Texas at Austin, TX. 200 mL of Expi293 cells suspension (GIBCO, Waltham, MA, USA) were transfected with the plasmid encoding the stabilized SARS-CoV-2 soluble S protein with the Expifectamine kit (GIBCO, Waltham, MA, USA) following manufacturer’s protocol. After six days of incubation, cell suspension was centrifuged, and supernatant clarified by 0.22 µm filtration. Supernatant was then processed through a His-tag cobalt resin (ThermoFisher, Waltham, MA, USA) following manufacturer’s protocol. The His-tag S protein was purified by gel filtration using the Akta explorer 100 system with a high-load Superdex pg 200 16/600 column (GE Life Sciences, USA). Main peak fraction was collected, and protein purity was verified by BN PAGE.

### Generation of ACE2+ 293T cells

The 293T cells were seeded in a 6-well plate and infected with a hACE2 lentivirus. Infected cells were incubated for 72 hours. After the incubation period cells were treated with 500ug/ml Hygromycin B (Fisher Scientific, Waltham, MA, USA) for about 14 days to select stable ACE2 expressing cells. Surviving cells were grown to confluency in a T75 flask and aliquots of cells were cryopreserved in complete DMEM/10%DMSO in liquid nitrogen.

### Pseudovirus and lentivirus production

Five million 293T cells were seeded in a T75 flask and let adhere overnight. The day after, the medium was replaced with 1:1 optiMEM (GIBCO, Waltham, MA) and 10% FBS DMEM (GIBCO, Waltham, MA, USA). 75ul Lipofectamine2000 (Invitrogen, Waltham, MA, USA) were added to 1.8ml optiMEM. To produce lentivirus carrying ACE2 expression cassette 14.2ug of pLENTI-ACE2 (ADDGENE, Watertown, MA, USA), 16ug of pCMVdR 8.2 (ADDGENE, Watertown, MA), and 6.4ug of pCMV-VSV-G (ADDGENE, Watertown, MA, USA) were mixed in 1.8ml optiMEM. To produce SARS-CoV-2 pseudotyped lentivirus 16ug of pCMDdR8.2, 14.2ug of pHAGE-Luc-zGFP (BEI resources, Manassas, VA, USA), and 6.4ug of CoV-2 glycoprotein expression plasmid (Wuhan version: pHDM BEI resources; for the BA.4/5 pLV-Spike V13 invivoGen plv-spike-v13) were mixed in 1.8ml complete optiMEM. Both Lipofectamine-optiMEM and Plasmid-optiMEM solutions were incubated 5 minutes RT. After incubation, plasmid solution and lipofectamine2000 were mixed and incubated for 15 minutes RT. After incubation, the final solution was added dropwise to the cells while gently shaking. The day after, the medium was replaced with 15ml complete DMEM and 1x non-essential amino acids (GIBCO, Waltham, MA, USA). Two supernatant batches were collected after 24 and 48 hours. The supernatant was collected and filtered with 0.45um PVDF Millex-GV filter (Millipore,Burlington, MA, USA). The cleared medium was supplemented with 1:5 Lenti Concentrator (OriGene, Rockville, MD, USA) and incubated 2-4 hours at 4°C. The cold solution was then centrifuged at 4000 RPM at 4°C for 40 minutes. Supernatant was removed and the pellet resuspended in 1/75 of the initial volume with complete DMEM. 70ul aliquots of the virus were stored at -80°C.

### Pseudovirus infection inhibition assay

The ACE2+ 293T cells or H838 cells were trypsinized and seeded in a 96-well plate at a concentration of 5000 cells per well in a volume of 100ul of DMEM complete medium. Cells were incubated and allowed to attach for 2 hours at 37C/5% CO2. In a separate plate, treatment curves (treatment concentration + complete DMEM) were prepared by serial dilution in a final volume of 50ul. After the incubation, the treatment was added to the cells and let incubate overnight. The treatments were with the following: *P. gingivalis*, *P. gingivalis* triple-gingipain knockout, or SPT knockout supernatants in 10%FBS DMEM, TSB, or BHI as well as *P. gingivalis* LPS or PGDHC compounds. Treatment with supernatants from bacteria grown from human saliva samples was also performed.

Pseudoviruses expressing the spike protein were added the next day. In another 96-well plate, the virus and treatments of either 293T or H838 cells were mixed in another serial dilution and left to incubate for 4 hours before 50ul were added to the cells in a final volume of 200ul. 72 hours after infection, luciferase was measured using BriteLite Plus (Perkinelmer, Waltham, MA, USA) and SynergyH1 plate reader.

### Postentry virus inhibition assay

To assess if our compounds have post-entry effect, the infection was carried out by spinoculation. In brief, the cells were exposed to the virus for 4 hours followed by 40 minutes centrifugation at 3000 RPM. The medium was removed, the cell washed with PBS and supplemented with fresh medium. The day after, the appropriate treatments were applied. The Luciferase activity was measured 72 hours after using BriteLite as described above.

### Syncytium formation assay

ACE2+ 293 were transiently transfected with pALPS expressing tat-p2a-rev cassette (Addgene, Watertown, MA, USA). The TZM.bl cell were transiently transfected with Wuhan CoV-2 spike expression plasmid pHDM (BEI resources, Manassas, VA, USA). ACE2+ 293 expressing tat/rev cells were treated with *P. gingivalis* supernatant, PGDHC, or medium alone overnight. The day after, TZM.bl expressing spike were added to the 293 ACE2+ expressing Tat/rev cells and incubated for 48 hours. After 2 days, the co-cultures were assessed for the formation of syncytia and luciferase activity as described above.

### WST-1 cytotoxicity assay

293 ACE2+ cells, at ∼90% confluency, were trypsinized and seeded in a 96-well plate at a concentration of 5,000 per well in a volume of 100ul of DMEM complete medium. Cells were incubated and allowed to attach for 2 hours at 37C at 5% CO2. In a separate plate, treatment curves were prepared by serial dilution in a final volume of 50ul per well. After the incubation, the treatment was added to the cells and let incubate for 72 hours. The treatments used are as follows: Pg Supernatant in TSB, BHI, and 10%FBS DMEM, Pg LPS, Pg RGPA/RGPB/KGP Knockout Supernatant in 10%FBS in DMEM, TSB, BHI, 10% FBS DMEM, PGDHC. On the third day toxicity was assessed by WST-1 (Cayman chemicals, Ann Arbor, MI, USA). The medium was removed and replaced with 100ul of our complete DMEM plus 10ul of WST-1. 450nm absorbance was then measured using the SynergyH1 plate reader (BioTek, Winooski, VT, USA) every 5 minutes over the course of two hours.

### Biolayer interferometry assay

To assess the ability of *P. gingivalis* supernatant and PGDHC to interfere with the ACE2-CoV-2 Spike protein interaction, we loaded anti-human IgG Fc bio sensors (Gatorbio, Palo Alto, CA, USA) for 360 seconds in a solution of 5ug/ml ACE2 Fc-tagged protein (ACRO Biosystems, Newark, DE, USA) in Buffer K (Gatorbio, Palo Alto, CA, USA)). After a 120-minute baseline, the biosensors were incubated for 300 seconds in a solution containing 15% *P. gingivalis* supernatant, 15ug/ml PGHDC or medium alone control. Following another 120-second baseline, the biosensors were immersed with a solution containing 10ug/ml of soluble CoV-2 Spike protein (see Supplementary Information) for 600 seconds. As last step, a 600 second dissociation step was performed in pure Buffer K.

We developed a method to analyze the inhibitory effect of PGDHC directly on pseudotyped lentiviruses. In brief, hFc biosensors (Gatorbio, Palo Alto, CA, USA) were loaded with fc tagged ACE2 protein (ACRO biosystems, Newark, DE, USA) until 1-1.5nm shift was reached. Following 120-second baseline, the sensors were dipped in a solution containing CoV-2 pseudotyped lentivirus, lentivirus with 5ug/ml neutralizing antibody, and virus with non-specific rabbit or non-specific IgG1/IgG2A antibodies.

## Statistical Analysis

Differences in viral infection susceptibility (measured by luciferase activity) was statistically evaluated using one-way ANOVA and Tukey post-hoc test for multiple comparison. T-test was used for single group comparison. The experiments were performed in triplicate from the same treatment preparations, and the means ± standard deviations were calculated. Differences with a probability of 5% (*P* < 0.05) were considered statistically significant.

## Data Availability

The datasets used and analyzed in this study are available from the corresponding author upon request.

## Acknowledgements

We thank Roodelyne Pierrelus, Sunniva Ruiz, and Margarita Uribe for assisting in saliva collection. We are grateful to Shin Nakamura for providing *P. gingivalis* cultures and supernatants. We thank Ms. Caitlin Dargue for excellent administrative support. This work was supported by the Florida Blue Foundation (M.C. and S.S.A.), NIH NIDCR R01-DE027249 (M.C. and A.B.), R01-DE031159 (M.E.D.), and NIGMS R16GM150469) (A.H.).

## Author contribution

Conceptualization, A.B., M.J.C, T.K., A.H.; methodology, A.B., A.C., M.J.C., C.G.G., A.L., M.R.P., F.J.N., J.P., M.E.D.; validation, A.B., M.J.C., A.H., C.G., A.C.; resources, M.J.C., T.K., S.S.A.; data curation, A.B., M.J.C., A.H.; writing/original draft preparation, A.B, M.J.C., A.C., T.K, M.E.D., F.C.H; supervision, A.B., M.J.C., T.K.; project administration, M.J.C., S.A.A., T.K., A.B., C.G. All authors have read and agreed to the published version of the manuscript.

